# Disentangling genetic variance for pathogen avoidance and resistance

**DOI:** 10.1101/2025.03.09.642266

**Authors:** Caroline R. Amoroso, Janis Antonovics

**Affiliations:** Department of Biology, University of Virginia, Charlottesville, VA, USA

## Abstract

Hosts can use avoidance (e.g., behavior) to reduce their contact rates with pathogens; after contact, they can use resistance (e.g., immunity) to reduce the establishment and proliferation of an infection. Because both defenses preserve host fitness and reduce pathogen fitness, we expect that their epidemiological and evolutionary effects will be inter-dependent. This study used a two-locus model to understand the evolution of allelic associations (i.e., linkage disequilibrium) between genes determining levels of avoidance and resistance in the presence of an infectious disease or parasite. We found that polymorphism in both avoidance and resistance was possible, but only for a limited range of parameter values of avoidance and resistance; at equilibrium, avoidance and resistance alleles were negatively associated in these polymorphic populations. However, most commonly, whichever defense was more effective and less costly went to fixation. This result suggests that avoidance and resistance may be more likely to covary negatively across than within populations.

## Introduction

Hosts are under selection to reduce the negative fitness effects of parasite infection. They can mitigate the negative effects of infection by *avoiding* initial contact with parasites, and/or by *resisting* parasite establishment and proliferation after contact has occurred [1]. Avoidance and resistance have shared effects: they increase host fitness in the presence of parasite risk, and they have negative effects on parasite fitness by limiting onward transmission [2]. Avoidance is commonly behavioral (synonymous with “behavioral resistance” in some studies), and resistance is commonly physiological (e.g., immune system) [3]. The goal of this study is to identify how the presence of one defense influences the evolution of the other using a basic two-locus model. Given their shared outcome, we expect that avoidance and resistance will interact epistatically, and thus that their evolutionary trajectories will not be independent.

In natural systems, evolution is predicted to shape patterns of co-occurrence of avoidance and resistance [4]. A negative correlation between avoidance and resistance phenotypes across individuals within the same population was found in several species of birds [5–7]. In salmon and trout, there was no relationship between avoidance and resistance to a parasitic eye fluke within populations [8], but a subsequent study found evidence of a negative relationship across populations [4]. On the other hand, the parasite *Serratia marcescens* caused the evolution of higher levels of both avoidance and resistance in *Caenorhabditis elegans* over 30 generations [9, 10]. These opposing examples highlight that we do not have a theoretical expectation for what drives the direction of covariance between avoidance and resistance, including whether their genetic basis plays a role, and at what scale we should expect to detect it [2].

Our understanding of resistance evolution has been strongly influenced by theory predicting that costs can help maintain genetic diversity in resistance [11, 12]. Empirical studies provide evidence that resistance to parasites can have fitness costs, and these costs may underlie genetic diversity in resistance [13–15].

On the other hand, although avoidance by behavior or phenological shifts has been described in a variety of plant and animal hosts, theoretical approaches to understanding avoidance evolution have been far fewer [16, 17]. We have shown previously that if avoidance costs were constitutive, polymorphism in avoidance behavior could be maintained, consistent with resistance theory. However, if avoidance costs varied as a function of the level of avoidance, it was less likely to evolve [16].

In this study, we examine how avoidance and resistance evolve when both defenses affect host fitness by building on previous models of avoidance evolution [16]. We ask: under what conditions do populations maintain both avoidance and resistance alleles, and in which direction do they associate? We examine how genetic variation in both traits is maintained and identify general conditions leading to positive or negative association of avoidance and resistance alleles.

### The Model

#### Definitions

Or model assumes an infectious agent (i.e., a pathogen or microparasite) that is directly transmitted between hosts that become infected. We define *defense* as a host trait that reduces the probability of becoming infected. In our model, the two possible defenses are avoidance and resistance. *Avoidance* reduces the probability of contact with infectious individuals, and *resistance* reduces the probability of becoming infected after contact. Another commonly studied defense is tolerance [18]; for the sake of simplicity, we assume that there is no variation in tolerance, and all infected individuals are equally infected and infectious. We assume avoidance and resistance are each controlled by a single genetic locus with two alleles. For both avoidance and resistance, we define *effectiveness* as the proportion by which an allele reduces the probability of infection, and the *cost* of an allele as the fraction by which it reduces the host’s birth rate (described in more detail below). We consider linkage (i.e., genetic distance) between the avoidance and resistance loci by varying their recombination rate. We examine linkage disequilibrium, (LD) or the degree of allelic association between the loci when the population has reached equilibrium (i.e., as an output of the model processes) [19]. We define the LD or allelic association as positive when the higher effectiveness (avoidance and resistance) alleles at both loci occur together more often in a genotype than expected based on their allele frequencies, or when the lower effectiveness (undefended) alleles at both loci occur together more often in a genotype than expected based on their allele frequencies. We define the LD or allelic association as negative when within one individual, alleles with higher effectiveness of one defense are more often associated with alleles with lower effectiveness of the other defense.

#### Model details

##### Host population demography and genetics

We use a Susceptible-Infected (SI) compartment model, where adult *S* individuals are susceptible and *I* are infected and infectious. For simplicity, we assume a population of well-mixed haploid hosts where host individuals vary at two loci, such that locus *A* controls avoidance of the parasite, and locus *B* controls resistance. At locus *A* the allele can be either *A*, which produces a more avoidant phenotype, or *a*, which produces a less avoidant phenotype. Similarly, at locus *B*, the allele is either *B*, which produces a more resistant phenotype, or *b*, which produces a less resistant phenotype. We refer to the genotype *ab* as the undefended genotype because it possesses neither defense. We use subscripts as in *S*_*ij*_ such that *i* and *j* are the allelic states for loci *A* and *B*, respectively. We assume that the pathogen is genetically invariant in its transmission and that once *S*_*ij*_ individuals become *I*, their infections are identical.

New births are denoted by 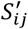, with the prime superscript to indicate that the number of genotype *S*_*ij*_ in the next generation includes recombination between parental genotypes, determined by the recombination rate, *R*, and random mating among the genotypes (Appendix 1). Offspring are produced at a baseline birth rate *b*, which is sensitive to density of the population according to 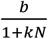, where *k* is a constant and *N* is the total population size. All hosts, including susceptible and infected, die according to a background mortality rate, *µ*.

Hosts have a fitness cost of defense *c*_*ij*_ that depends on their genotype such that their birth rate is reduced by (1 −*c*_*i*_)(1 −*c*_*j*_). The cost of the *A* allele is *c*_*A*_, and the cost of the *B* allele is *c*_*B*_. In the *AB* genotype, the cost is multiplied, such that the two loci have an epistatic effect on birth rate. We assume *c*_*a*_ and *c*_*b*_ are both 0, so the undefended genotype has a birthrate equal to *b*.

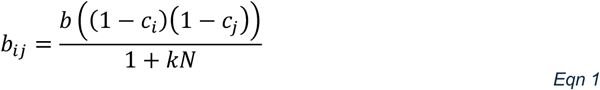

##### Pathogen infection dynamics

The transmission rate (*β*) is determined by the host genotype, *β*_*ij*_ and the frequency of infected individuals in the population, or prevalence, 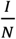. This assumption reflects frequency-dependent transmission, which we chose for simplicity (*β*_*ij*_ varies from 0 to 1) and to conform to prior models [11, 16].

Our model produces the following generalized differential equations:

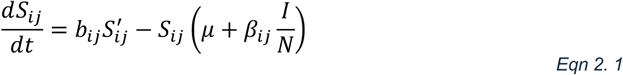

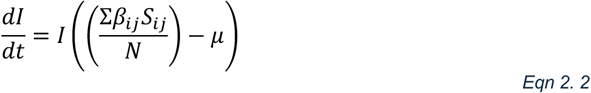

Avoidance and resistance at loci *A* and *B* respectively reduce transmission. The magnitude of these defenses is represented by *e*_*A*_, the effectiveness of avoidance, and *e*_*B*_, the effectiveness of resistance. This reduced transmission probability is relative to the undefended genotype, *S*_*ab*_; *e*_*a*_ and *e*_*b*_ = 0, so the undefended transmission probability (*β*_*AB*_) is equal to δ. For *S*_*AB*_, which has both defenses, the defenses’ effects are multiplied to reflect that avoidance reduces the proportion of contacts that the host makes, and resistance reduces the proportion of the remaining contacts that proceed to infection. This produces an epistatic effect between the two defense traits on transmission probability:

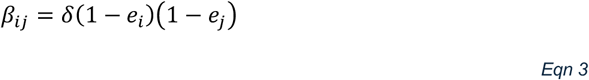

For consistency with previous theory, we assume the infection is sterilizing. The parameters of the model are summarized in Table 1 with the values used for those parameters in the simulations (unless otherwise indicated). The full equations are in the Supplementary Materials (Equation S1.1-S1.5).

**Table 1.** Parameter descriptions and the values used in the simulations.

| Parameter | Description | Values |
| --- | --- | --- |
| $b$ | Baseline birth rate | 1 |
| $R$ | Recombination rate | 0.01, 0.5 |
| $k$ | Degree of density dependence | $10^{-5}$ |
| $\mu$ | Background mortality rate | 0.1 |
| $\delta$ | Baseline pathogen transmission rate | 0.8 |
| $e_A, e_B$ | Effectiveness of defense of avoidance (the $A$ allele) or resistance (the $B$ allele), respectively | (0,1) |
| $c_A, c_B$ | Constitutive cost of possessing avoidance (the $A$ allele) or resistance (the $B$ allele), respectively | (0,1) |

##### Simulations

We simulated populations that were governed by the above differential equations programmed in R (version 4.4.1) [20] using the ordinary differential equation solver from the package *deSolve* (function ‘ode,’ classical 4^th^ order Runge-Kutta method, i.e., *method= “rk4)* [21]. We used the *tidyverse* package for visualization [22]. All simulations were deterministic. We simulated our model over 2000 timesteps. To confirm that the equilibria were non-cyclic, we perturbed values above and below equilibrium values to confirm their stability. Note that after 2000 timesteps, some of the bordering regions that appear as narrow strips of parameter space producing a condition in the plotted results may be better represented by a line if they were more fully resolved over a greater number of timesteps.

Previous models of a single locus with two alleles of avoidance [16] and resistance [11, 23] showed three possible equilibrium states: the defended (avoidance or resistance) allele could become fixed, the undefended allele could become fixed, or the two alleles could coexist at intermediate frequencies in a stable polymorphism. Our model confirmed these results when we only examined variation in one defense at a time. From this parameter space, we selected a representative point from each of these three regions (referred to as Points I, II and III). To examine how avoidance evolved under each of these three assumptions about resistance costs and effectiveness, we introduced variation in avoidance at the second locus. We focused on two different assumptions about how this variation in avoidance was introduced. We first examined how evolution would proceed from standing genetic variation at both loci, i.e., starting with a population that had all four genotypes present at equal frequencies and with no LD. As a follow-up, we also examined how evolution would proceed if avoidance was introduced at a low frequency (e.g. by mutation or migration) into populations that had reached a stable equilibrium in resistance.

##### Avoidance evolution from standing variation in both defenses

To simulate evolution from standing genetic variation, we started our simulations with all four genotypes present at equal frequencies and therefore at linkage equilibrium. We ran simulations with the effectiveness and cost of avoidance set to all the combinations of values within the range of 0 to 1, and other parameters at base values. For recombination rate (*R*), we ran the simulations with values of 0.01 (low recombination; highly linked), and 0.5 (free recombination; completely unlinked). (We also explored the specific case of complete genetic linkage, *R* = 0, and show results in the Supplementary Materials.)

##### Avoidance invasion

To explore how initial starting frequency and linkage disequilibrium affected the outcome when the avoidance allele was introduced at a low frequency, we selected the same three representative points from the resistance-only simulations. Then we added variation in avoidance by introducing only either genotype *Ab* (Points I and III) or *AB* (Points II and III) at a frequency of 10^−5^ of the equilibrium population size. Then we continued the simulation for another 2000 timesteps. We again ran simulations with the effectiveness and cost of avoidance over the range of 0 to 1, and other parameters at base values and recombination as previously. These results are described briefly in the main text and presented in full in the Supplementary Materials (see Results for reference to specific sections).

## Results

In our model, avoidance and resistance could each independently reduce transmission relative to the undefended genotype, and they reduced it maximally when they were both present (Fig. S1 in the Supplementary Materials). Thus, in the absence of costs, the genotype with both defenses would increase in frequency and become fixed in the population. Varying the cost and effectiveness of resistance produced outcomes consistent with previous models of defense evolution. When the effectiveness of resistance was low, it was unlikely to evolve (*ab* reached fixation). As resistance increased, it could evolve at a greater cost (*aB* reached fixation). At high effectiveness of resistance and across a wide range of costs, there was a stable coexistence between the resistance allele *B* and the less defended allele *b*. In this region, the more resistant allele lowered the prevalence of infection, enabling the undefended, less costly genotype to persist at equilibrium (Fig. 1). The results presented here focus on the dynamics when a population reaches equilibrium; however, this equilibrium state does not necessarily reflect the transitory patterns as the population evolved, prior to reaching equilibrium. For example, a population in which one allele reaches fixation at equilibrium (producing a value of zero for linkage disequilibrium) may have experienced periods of positive and negative linkage before equilibrium (Fig. S2 in the Supplementary Materials). The direction of the initial linkage in some cases depends on the infection prevalence.

**Figure 1.**
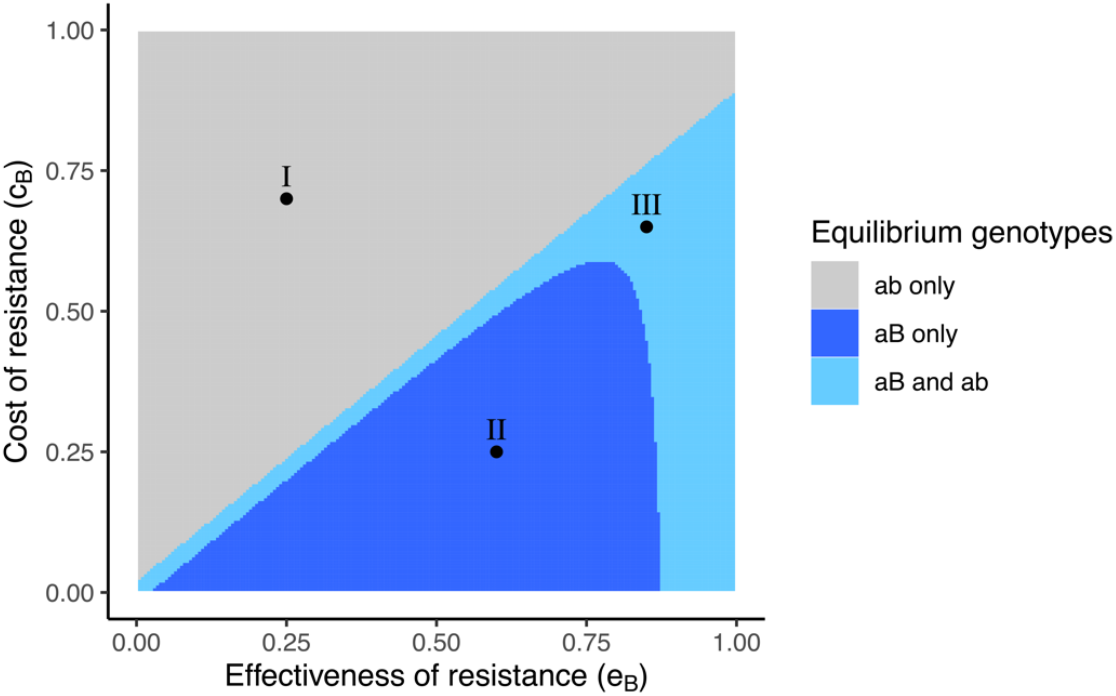
The effectiveness and cost of resistance influence its evolution. Regions of presence for different genotypes at equilibrium when the effectiveness (*e*_*B*_) and cost (*c*_*B*_) of resistance are varied, assuming there is no avoidance and only considering variation in resistance. Three representative points from within this parameter space were selected for further exploration (labeled I, II, and III). Aside from *c*_*B*_ and *e*_*B*_, the parameters were set as follows: *b* = 1, *k* = 10^−5^, *µ* = 0.1, *δ* = 0.8.

### Avoidance evolution from standing variation in both defenses

To examine how avoidance evolution varied across these resistance conditions, we selected a point from each of the three equilibrium outcome regions (labeled I, II, and III in Fig. 1) at which to fix the effectiveness and cost of resistance.

We first examined how avoidance evolved when the population was fixed for allele *B* (i.e., Point I in Fig. 1: *e*_*B*_ = 0.25, c_*B*_ = 0.7). Avoidance (*Ab*) evolved at all levels of effectiveness when its costs were low, and at increasing costs as effectiveness increased. There was also a region of coexistence between *ab* and *Ab* at moderate to high effectiveness across a range of avoidance costs. As expected, varying the recombination rate did not change the outcomes, because at Point I, the population was fixed for *b* (Fig. 2).

**Figure 2.**
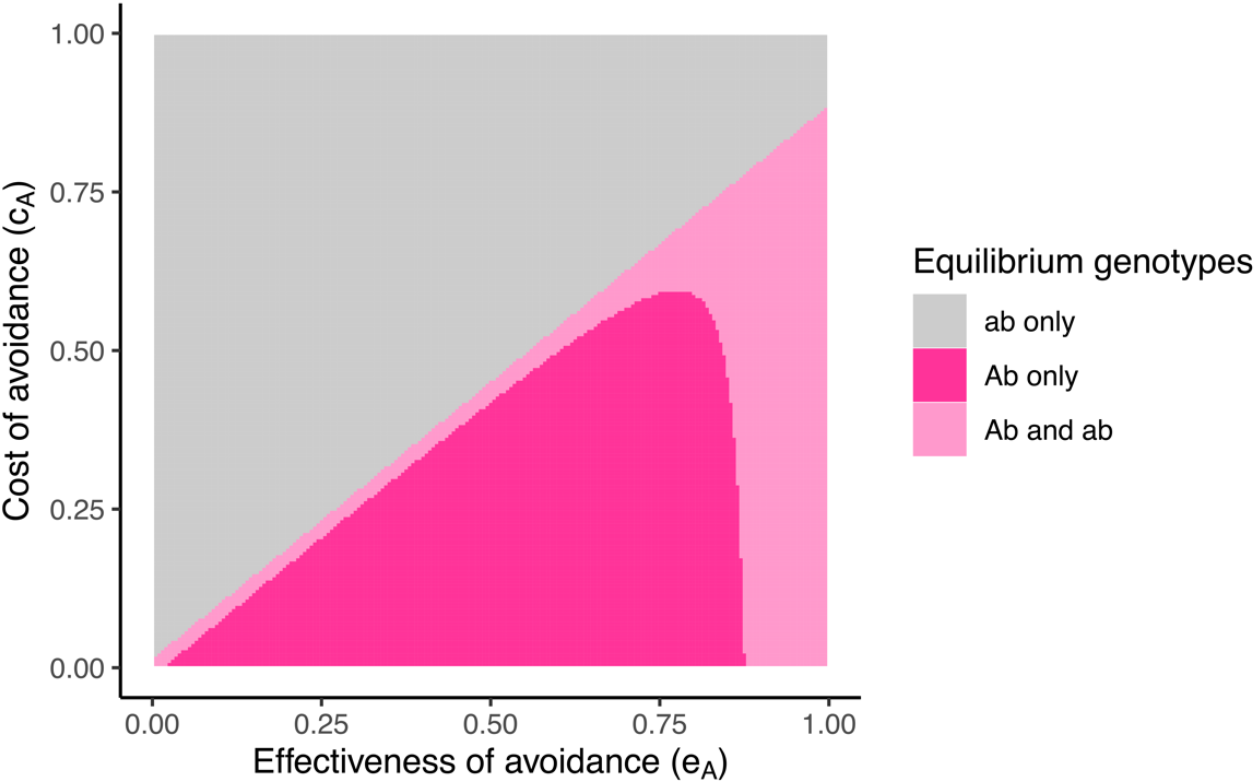
Evolution of avoidance from standing variation at both loci under conditions that did not support resistance evolution. Regions of presence for different genotypes at equilibrium when the effectiveness (*e*_*A*_) and cost (*c*_*A*_) of avoidance were varied. The cost and effectiveness of resistance were held constant at the values represented by Point I in Figure 1: *r*_*B*_ = 0.25, *c*_*B*_ = 0.7, at which *b* became fixed when there was no variation in avoidance.

Next, we examined conditions for avoidance evolution when resistance (*B*) had gone to fixation for b, having a low cost (Point II in Fig. 1, *e*_*B*_ = 0.6, *c*_*B*_ = 0.25). When avoidance costs were high, increased avoidance did not evolve, leaving resistance as the lone defense (Fig. 3, upper left half of the plot). At low avoidance and low cost, *AB* evolved to fixation (lower left) or in polymorphism with *Ab* or *aB* (lower central, and central-diagonal regions, respectively). At higher levels of avoidance effectiveness (i.e. *e*_*A*_ exceeding the fixed *e*_*B*_ = 0.6 at Point II) and lower costs, avoidance was able to supplant pure resistance (*aB*), with *Ab* evolving to fixation or in polymorphism with *AB* or *ab*. At the greatest levels of effectiveness and intermediate costs of avoidance, the outcome depended on the recombination rate between the two loci. At low levels of recombination, all four genotypes were maintained in a polymorphism at high levels of avoidance effectiveness and intermediate costs (green area of Fig. 3a). In this region, there was negative LD or allelic association between the levels of avoidance and resistance (Supplementary Figure S2) However, when there was free recombination, all four genotypes could only be maintained in a very small region of parameter space where the costs and effectiveness of avoidance were equal or close to the values of the parameters for resistance (Fig 3b: *e*_*A*_ ≈ *e*_*B*_ = 0.6 and *c*_*A*_ ≈ *c*_*B*_ = 0.25).

**Figure 3.**
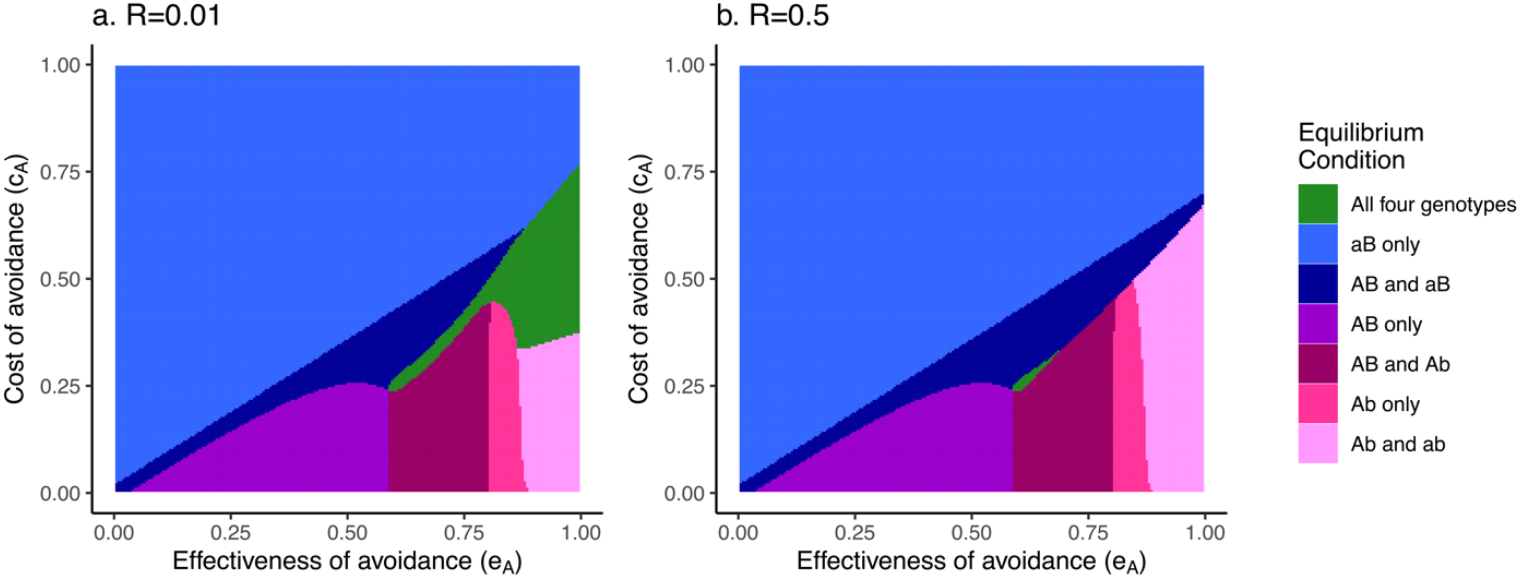
Evolution of avoidance from standing variation at both loci under conditions that supported resistance evolution. Regions of presence for different genotypes at equilibrium when varying the cost (*c*_*A*_) and effectiveness (*e*_*A*_) of avoidance and holding the cost and effectiveness of resistance constant at the values represented by Point II in Figure 1: (*e*_*B*_ = 0.25, *c*_*B*_ = 0.7), at which *B* became fixed when there was no variation in avoidance. Outcomes of simulations assuming low (a) and high (b) recombination rates (*R*) are shown in the panels.

In the region where *ab* and *aB* were polymorphic in the absence of avoidance (Point III in Fig. 2, *e*_*B*_ = 0.85, *c*_*B*_ = 0.65), introducing variation at the avoidance locus produced a region where the genotype *Ab* evolved in the area where avoidance effectiveness was high and costs were low (Fig 4, lower right area of the plots). At lower effectiveness and costs, *Ab* was polymorphic with *AB*, and at higher effectiveness, *Ab* was polymorphic with *ab*. When there was a low recombination rate, it was possible for all four genotypes to be maintained across a narrow diagonal strip of parameter space (green region in Fig. 4a), and in this area, avoidance and resistance were negatively associated (Supplementary Figure S3). The variation in both defenses was no longer maintained in this region when there was free recombination, other than at the very lowest levels of cost and effectiveness (green region in Fig. 4b).

**Figure 4.**
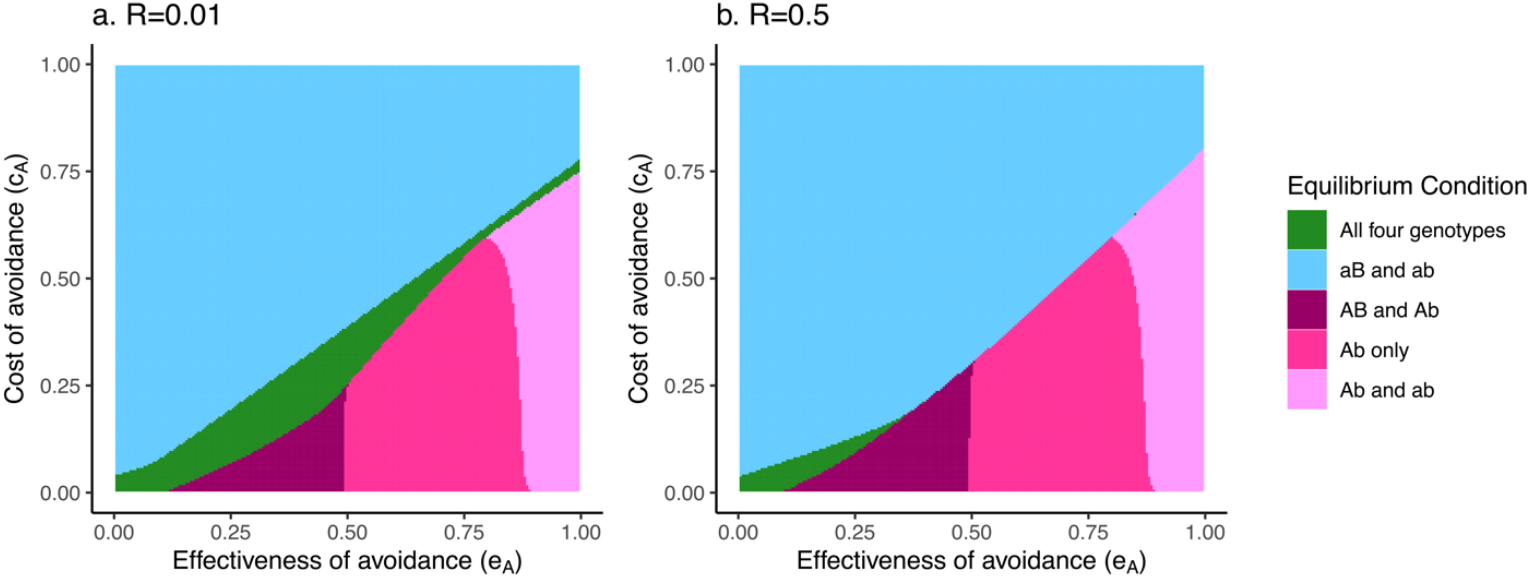
Evolution of avoidance from standing variation at both loci under conditions that supported resistance polymorphism. Regions of presence for different genotypes at equilibrium when varying the cost (*c*_*A*_) and effectiveness (*e*_*A*_) of avoidance and holding the cost and effectiveness of resistance constant at the values represented by Point III in Figure 1: *e*_*B*_ = 0.85, *c*_*B*_ = 0.65. In the absence of avoidance variation, these parameters led to stable polymorphism in resistance. Outcomes of simulations assuming low (a) and high (b) recombination rates (*R*) are shown in the different panels.

To summarize, when a population has standing genetic variation at both loci, there are specific conditions that allow this variation in both avoidance and resistance to be maintained (i.e., the green regions in Figs. 3 and 4); however, over a large range of parameters, it is much more common for variation in at least one defense to be lost.

### Avoidance invasion at low frequency

We also considered whether avoidance could evolve when introduced at a low frequency, for example, if avoidance variation were the result of a novel mutation. As long as there was some amount of recombination possible, the equilibria produced through invasion mirrored the results for standing genetic variation (Supplementary Figs. S5-S7). One case that differed, however, was when resistance was held at the values for Point II (fixed for *S*_*aB*_) and avoidance was introduced at a low frequency in positive association with resistance (as *S*_*AB*_). In these cases, the population evolved to *AB* only, or to *AB* and *aB* (Supplementary Figure S7). In the special case where the two loci were completely linked (*R* =0), the equilibrium state depended on whether avoidance was introduced in positive or negative association with resistance (as *AB* or as *Ab*) (Supplementary Figs. S8 and S9).

## Discussion

We have described here simulations of a population-genetics model that considers the evolution of both avoidance and resistance to a directly transmitted pathogen. A central conclusion from our model is that because of their shared outcome, assumptions about the cost and effectiveness of one defense can have a large impact on the evolution of another. Whether a population already has variation in resistance has a large effect on when avoidance evolution is favored. Assumptions about the linkage relationships (i.e., genetic distance) between these defenses also have a strong bearing on evolutionary outcomes. It is worth noting that that our results focus on the equilibrium states of the modeled populations, which addresses expectations for a long and stable host-pathogen relationship. In the shorter term, when a pathogen is first introduced into a host population, or when a defense is first introduced, there can be transient associations between avoidance and resistance that emerge briefly and even change direction (e.g., Fig. S2). The trajectory of these associations is not necessarily reflected by the equilibrium state, and depends not only on the relative cost and effectiveness of the two defenses, but also on the infection prevalence.

One surprising finding was how rarely we observed the evolution of populations that would be considered to demonstrate a “trade-off” between avoidance and resistance within a population. Within a population, this implies there would be individuals with both high avoidance but low resistance, and many with the opposite [5, 6, 8]. The results from the current study contrast with this expectation, especially if many combinations of costs and effectiveness of the two defenses are possible. Across most of the model parameters, variation in at least one defense was lost when the population reached equilibrium. When populations did maintain variation at both loci, avoidance and resistance alleles were indeed negatively associated (i.e., in negative LD).

However, these regions were small, such that it might be expected that slight perturbations to model parameters, like the level of density dependence or background mortality, to nudge the population out of equilibrium and cause one or the other defense to predominate. Our results suggest that although a trade-off between avoidance and resistance is intuitively compelling as a verbal argument, a negative covariance would require quite specific conditions to be maintained within a population, at least under the assumptions of this model.

Instead, it was more common for one or the other defense to go to fixation, or for one or the other defense to be in polymorphism with the undefended genotype. This general pattern suggests that if the effectiveness and costs of avoidance and resistance varied across environments, and assuming the costs were not too high, most individuals within a population would evolve one defense or the other. If the costs and effectiveness of each defense - or any other model parameter - were different across environments, this could generate a negative covariance *across* populations, i.e., with some populations evolving primarily avoidance and others evolving primarily resistance. This pattern of negative covariance between avoidance and resistance across host populations has been described in at least one study in the literature [4]. We also observe avoidance and resistance evolving jointly (i.e., *AB*) when they were physically unlinked across a range of relatively low costs and low to moderate effectiveness. This positive covariance between avoidance and resistance also has some empirical evidence in the literature, especially in a system where costs of defense have not been detected, and so presumably are low [24, 25]. In general, this study raises the importance of the scale at which defense covariances are measured [4, 26].

As with most theoretical models, we have made simplifying assumptions in this study. We acknowledge that these simplifications have made the model more abstract, but in doing so, we uncover general patterns that may be broadly applicable to many different host-pathogen systems. These assumptions also point out gaps in our current knowledge about avoidance and resistance evolution, and we outline these below.

First, we model the transmission-reducing effectiveness of avoidance and resistance as simply multiplicative. However, the simple multiplication of these probabilities also leaves the order of that sequence undefined. Our model therefore treats avoidance and resistance as symmetrical, working in the same way to reduce transmission (as supported by the similarity of Figures 1 and 2). However, definitionally, avoidance precedes resistance, and the defenses work in a step-wise manner that could affect the evolutionary dynamics between them, with earlier defenses potentially blocking or shielding later defenses from selection [27, 28]. Avoidance defenses commonly take a behavioral form, while resistance defenses are commonly physiological [3] and the form of the defense may define how it reduces transmission risk, as well as how the effectiveness of a defense is associated to its costs. In any particular host-pathogen system, the broad spectrum of effectiveness and costs modeled here may not be possible, or may take different forms, changing the expectations for the defenses’ covariance.

Second, we assume the costs of avoidance and resistance are constitutive, such that a defense reduces fitness by a fixed amount [29]. This may be unlikely for avoidance if it takes a behavioral form and its costs result from using the defenses in response to disease incidence [3]. Future research could consider how modeling avoidance and its costs as induced by the presence of infection risk changes patterns of linkage disequilibrium.

Our results re-emphasize a broader point about the fact that epistasis can occur at two levels. The first is “trait epistasis” [30]. There is trait epistasis between the defenses in their effects on transmission because the allele present at one locus determines the magnitude of the effect at a second locus. There is also trait epistasis in the fecundity costs for the same reason. The second is “fitness epistasis,” which occurs between two loci anytime the relative fitness of the individual possessing two alleles of interest is not the exact product of the relative fitnesses of individuals each possessing one of the alleles [31]. Examining the full equations (Supplementary Materials Eqns S1.1-S1.4), it is clear that the relative fitness of *S*_*AB*_ is not the product of the relative fitness of *S*_*Ab*_ and *S*_*aB*_; therefore, avoidance and resistance are not only in trait epistasis but also in fitness epistasis, and expected to evolve linkage disequilibrium because of the latter [31, 32]. In our model, removing the trait epistasis by making the interactions additive does not remove the fitness epistasis. Indeed, it is difficult to imagine a theoretical model based in a susceptible-infected transmission framework where two loci would not be in fitness epistasis. Therefore, it seems inevitable that avoidance and resistance should influence each other’s evolution, at least to the degree that we accept this prevailing modeling framework.

A third simplifying assumption of this model is that there is no pathogen evolution. We are thus ignoring that coevolutionary dynamics between host and pathogen might act quite differently for the traits of avoidance and resistance. Previous studies have found that more flexible or plastic forms of defense (as predicted for avoidance) may select for different pathogen traits (e.g., lower virulence) than fixed defenses [33]. Accounting for these coevolutionary relationships could have a large impact on our model’s expectations. For example, if a model included variation in the pathogen population, questions would arise about whether the host defenses were equally effective against all variants of the pathogen, or whether their specificity differed. Avoidance is commonly predicted to be less specific than resistance [3], but there is recent evidence that avoidance in some hosts may be specific to pathogen genotype [34]. These factors could affect our evolutionary expectations, since the evolution of specific resistance can select against general resistance [35].

In conclusion, our model shows that avoidance and resistance are likely to influence each other’s evolution because of their joint effects on infection, and because of their potential costs. Even with the simplified nature of our assumptions, the model demonstrates that predictions about the relationship between avoidance and resistance are not necessarily straightforward or intuitive. The outcomes depend on how much protection from infection each defense provides, each defense’s costs, their interaction, and the ensuing ecological feedbacks on pathogen abundance.

