## Supplementary Materials for "Disentangling genetic variance for pathogen avoidance and resistance"

**Electronic Supplementary Material accompanying the following manuscript:**

**Supplementary Table 1: Mating matrix.** This matrix is used to determine how recombination contributes new offspring of each genotype to the next generation. $R$ is the rate of recombination, which varies from 0 (no recombination; loci are physically linked) to 0.5 (free recombination, i.e., independent assortment of loci). The proportion of new births comprised of each genotype is determined by multiplying each cell of column 2 by the frequencies in column 3 for the current timestep and taking the sum of the columns. Each proportion is then multiplied by the total number of adult $S_{ij}$ individuals to give values for $S_{ij}'$ in Eqn 2.1 and Eqns S1.1-S1.4.

| 1. Pairing | 2. Offspring genotype proportions | | | | 3. Frequency of pairing |
|---|---|---|---|---|---|
| | 2a. $AB$ | 2b. $Ab$ | 2c. $aB$ | 2d. $ab$ | |
| AB x AB | $1$ | 0 | 0 | 0 | $\left(\frac{S_{AB}}{N}\right)^2$ |
| AB x Ab | $0.5$ | $0.5$ | 0 | 0 | $2\left(\frac{S_{AB}}{N}\right)\left(\frac{S_{Ab}}{N}\right)$ |
| AB x aB | $0.5$ | 0 | $0.5$ | 0 | $2\left(\frac{S_{AB}}{N}\right)\left(\frac{S_{aB}}{N}\right)$ |
| AB x ab | $0.5(1-R)$ | $0.5R$ | $0.5(1-R)$ | $0.5R$ | $2\left(\frac{S_{AB}}{N}\right)\left(\frac{S_{ab}}{N}\right)$ |
| Ab x Ab | 0 | $1$ | 0 | 0 | $\left(\frac{S_{Ab}}{N}\right)^2$ |
| Ab x aB | $0.5R$ | $0.5(1-R)$ | $0.5R$ | $0.5(1-R)$ | $2\left(\frac{S_{Ab}}{N}\right)\left(\frac{S_{aB}}{N}\right)$ |
| Ab x ab | 0 | $0.5$ | 0 | $0.5$ | $2\left(\frac{S_{Ab}}{N}\right)\left(\frac{S_{ab}}{N}\right)$ |
| aB x aB | 0 | 0 | $1$ | 0 | $\left(\frac{S_{aB}}{N}\right)^2$ |
| aB x ab | 0 | 0 | $0.5$ | $0.5$ | $2\left(\frac{S_{aB}}{N}\right)\left(\frac{S_{ab}}{N}\right)$ |
| ab x ab | 0 | 0 | 0 | $1$ | $\left(\frac{S_{ab}}{N}\right)^2$ |

**Supplementary Equations : Full model**

$$\frac{dS_{ab}}{dt} = \frac{bS'_{ab}}{1+kN} - S_{ab}\left(\mu + \delta\frac{I}{N}\right)$$

*Eqn S1.1*

$$\frac{dS_{Ab}}{dt} = \frac{b(1-c_A)S'_{Ab}}{1+kN} - S_{Ab}\left(\mu + \delta(1-e_A)\frac{I}{N}\right)$$

*Eqn S1.2*

$$\frac{dS_{aB}}{dt} = \frac{b(1-c_B)S'_{aB}}{1+kN} - S_{aB}\left(\mu + \delta(1-e_B)\frac{I}{N}\right)$$

*Eqn S1.3*

$$\frac{dS_{AB}}{dt} = \frac{b\big((1-c_A)(1-c_B)\big)S'_{AB}}{1+kN} - S_{AB}\left(\mu + \delta(1-e_A)(1-e_B)\frac{I}{N}\right)$$

*Eqn S1.4*

$$\frac{dI}{dt} = \delta\frac{I}{N}(S_{ab} + (1-e_A)S_{Ab} + (1-e_B)S_{aB} + (1-e_A)(1-e_B)S_{AB}) - \mu I$$

*Eqn S1.5*

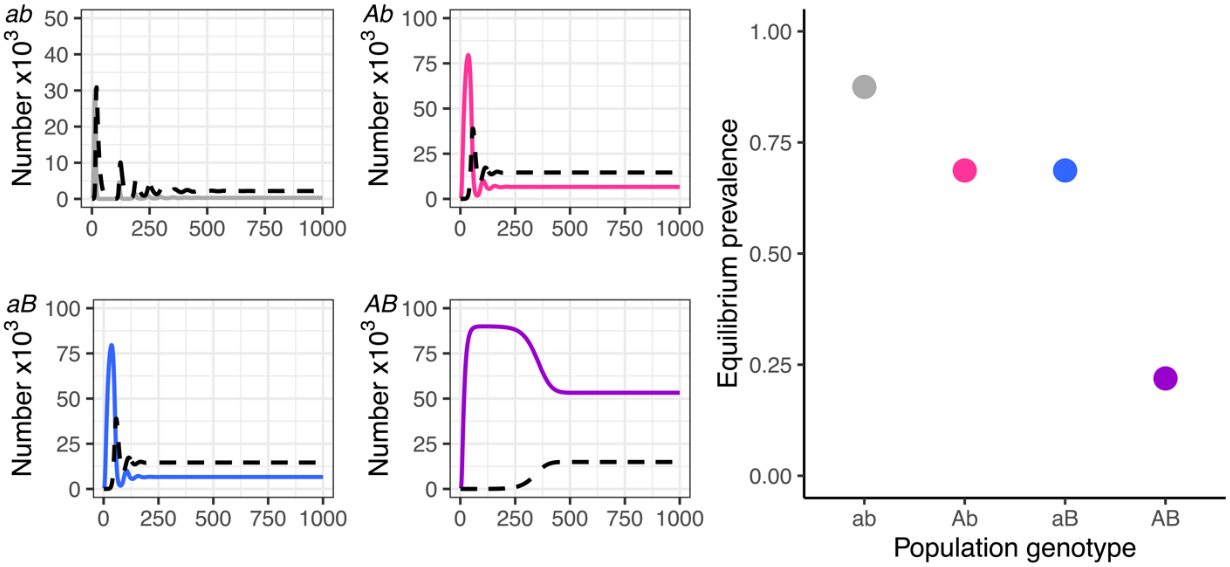


**Figure S1. Example simulations of single-genotype populations.** To illustrate the effects of avoidance and resistance independently and jointly in terms of reducing transmission, we first ran a version of the model with no variation in defense. This allowed us to examine transmission dynamics across populations that varied in their defense phenotype, in the absence of evolution. We assumed no cost of defense for the sake of illustration (i.e., $c_A = 0, c_B = 0$). The small panels display the of numbers of individuals in example simulations with only one genotype in each population across 1000 timesteps of the model. The black dotted line represents the number of $I$ (infected) individuals, and the solid lines represent the number of $S$ (susceptible) individuals of a given genotype. Each panel represents a population comprised of a single genotype without any costs of defense. The righthand scatterplot summarizes the prevalence at equilibrium in a population comprised of only that genotype; i.e., the proportion of infected at the 1000$^{th}$ timestep in the lefthand plots. The model parameters were held constant across the four simulations: $b = 1, c_A = 0, c_B = 0, k = 10^{-5}, \mu = 0.1, \delta = 0.8, e_A = 0.6, e_B = 0.6$. Note the scale of the y-axis for genotype $ab$ (upper left) is different than the others. From this plot, it can be seen that each defense independently reduced prevalence, and the genotype with both defenses had the lowest prevalence.

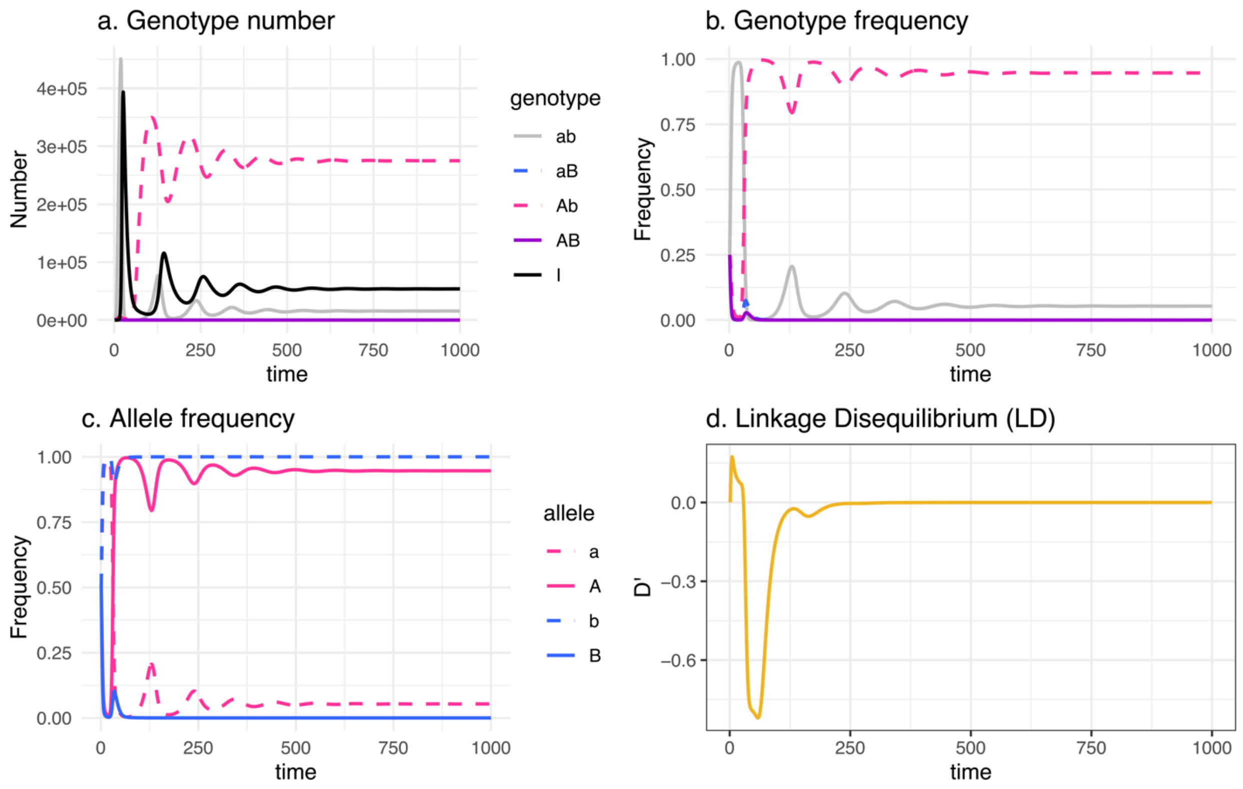


**Figure S2. Example simulation with all four genotypes present.** To demonstrate the distinction between equilibrium and transitory dynamics, we provide this example simulation across 1000 timesteps, including panels detailing changes over time in: a) the number of susceptible individuals of each genotype and the number of infected individuals (I); b) the frequency of each genotype out of the total susceptible individuals; c) the frequency of each allele; and d) a measure of LD, linkage disequilibrium, that is adjusted by the maximum possible value of D ($D_{max}$). At the start of the simulation, all four genotypes were present at equal frequencies, $N = 1000$, with a low prevalence ($\frac{I}{N} = 10^{-4}$). The other parameters were as follows: $b = 1, c_A = 0.5, c_B = 0.65, k = 10^{-5}, \mu = 0.1, \delta = 0.8, e_A = 0.9, e_B = 0.85, R = 0.01$. The resistance parameters correspond to Point III in Fig. 1, and the equilibrium state is a polymorphism between $Ab$ and $ab$. Note that the linkage disequilibrium ($D'$) reached 0 at equilibrium; however, it was initially positive when the infection prevalence was low and genotype $ab$ increased in frequency. Then, as infection prevalence increased, the avoidance allele, which was more effective and less costly than the resistance allele, increased in frequency as the genotype $Ab$ was selected, corresponding to a negative LD. As the $B$ allele decreased in frequency and $b$ reached fixation, LD approached and then equaled zero. This example illustrates that even if there were no association between alleles (i.e., LD=0) at equilibrium, there were likely to be non-zero associations between alleles that could even be in opposite directions. The direction of initial disequilibrium also depends on the starting prevalence, with low prevalences producing an initial positive LD, but high starting prevalences producing negative LD initially.

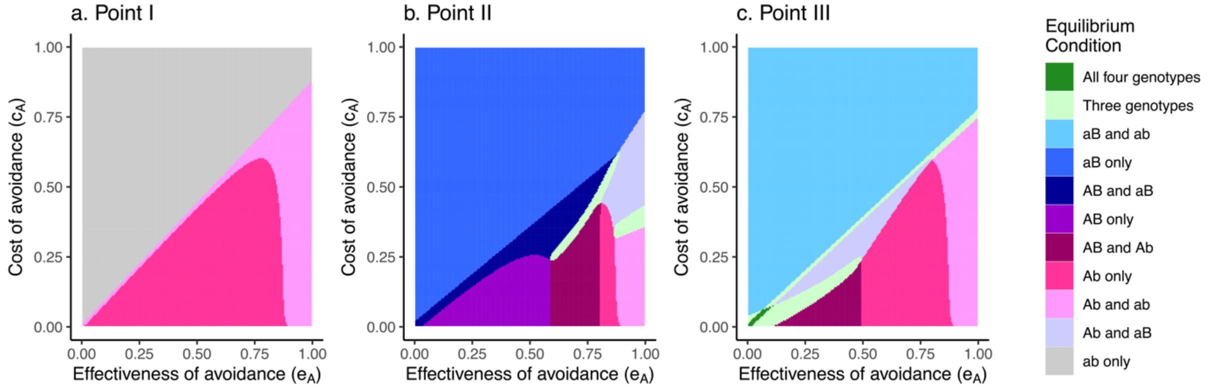


**Figure S3. Evolution from standing variation at both loci when avoidance and resistance loci are completely linked (R=0).** Regions of presence for different genotypes at equilibrium when varying the cost ($c_A$) and effectiveness ($e_A$) of avoidance and holding the cost and effectiveness of resistance constant at the values represented by Point $\mathrm{III}$ in Figure 1: $e_B = 0.85, c_B = 0.65$. In the absence of avoidance variation, these parameters led to stable polymorphism in resistance. Outcomes of simulations assuming low (a) and high (b) recombination rates ($R$) are shown in the different panels.

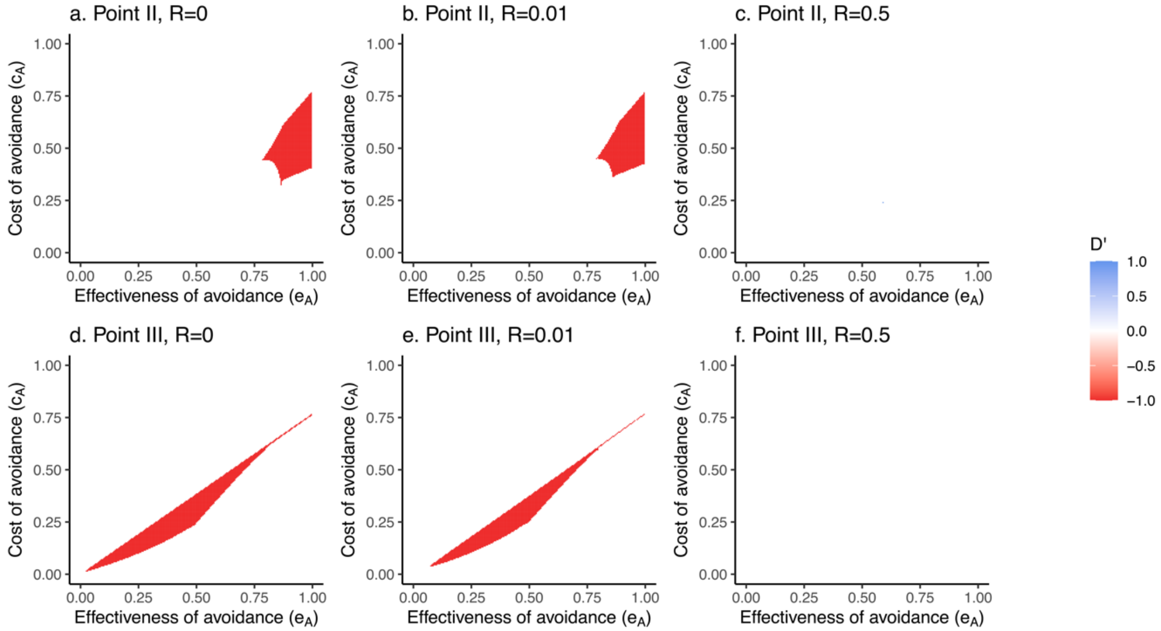


**Figure S4. Linkage disequilibrium (LD) at the populations' equilibrium states**. $D'$, linkage disequilibrium corrected for $D_{max}$ (per Lewontin 1988, ref. 19 in the main text), when varying the cost ($c_A$) and effectiveness ($e_A$) of avoidance and holding the cost and effectiveness of resistance constant at the values represented by Points $\mathrm{II}$ and $\mathrm{III}$ in Figure 1. $D'$ is zero for all populations at equilibrium when the cost and effectiveness of resistance are held constant at Point $\mathrm{I}$.

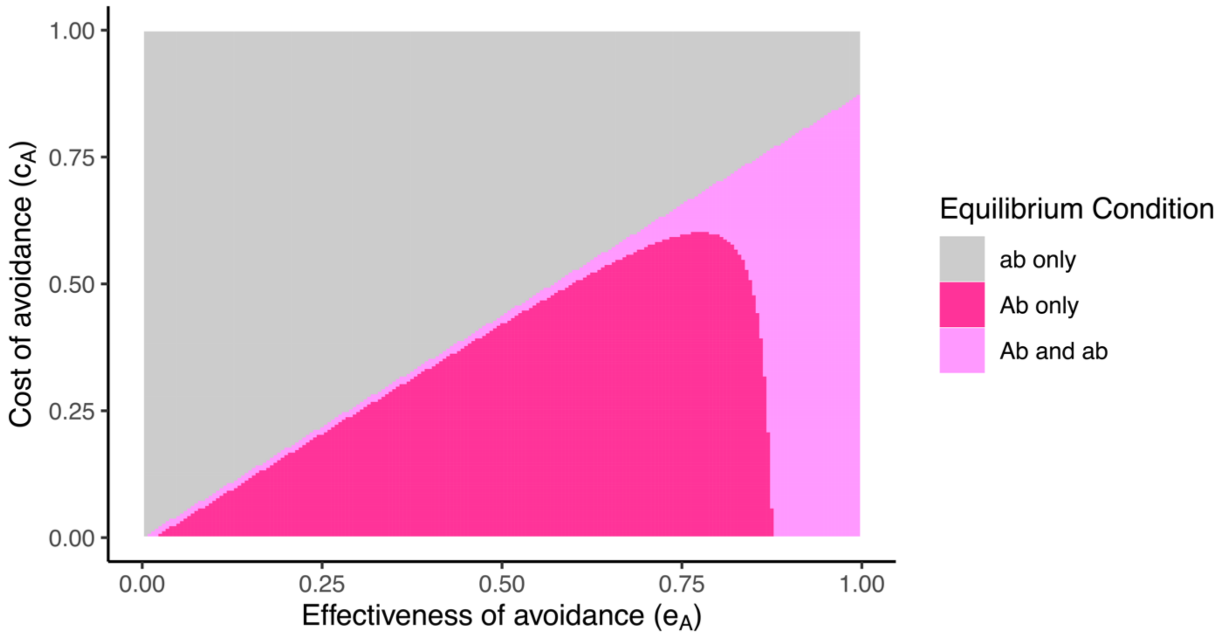


**Figure S5. Evolution from a low frequency at both loci under conditions that did not support resistance evolution (Point I).** The cost and effectiveness of resistance were set at the values represented by Point I in Figure 1: $r_B = 0.25, c_B = 0.7$, and run for 2000 timesteps, conditions leading $b$ to become fixed when there was no variation in avoidance. Then $S_{Ab}$ was introduced at a low frequency $(10^{-5})$ and the simulations were continued for 2000 timesteps. Setting the recombination rate $(R)$ to different values (0, 0.01, and 0.5) did not change the result.

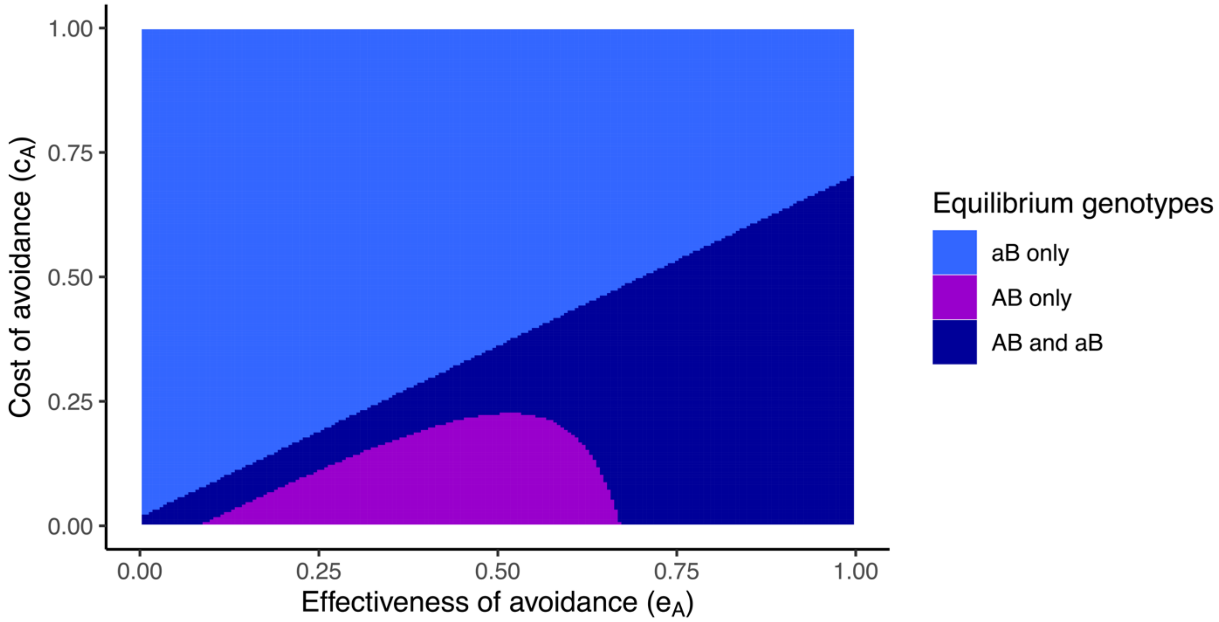


**Figure S5. Evolution of avoidance from a low frequency at both loci under conditions that supported resistance evolution (Point II).** The cost and effectiveness of resistance were set at the values represented by Point II in Figure 1: ($e_B = 0.25, c_B = 0.7$), and run for 2000 timesteps, under which conditions $B$ became fixed when there was no variation in avoidance. Then $S_{AB}$ was introduced at a low frequency ($10^{-5}$) and the simulations were continued for 2000 timesteps. Setting the recombination rate ($R$) to different values (0, 0.01, and 0.5) did not change the result.

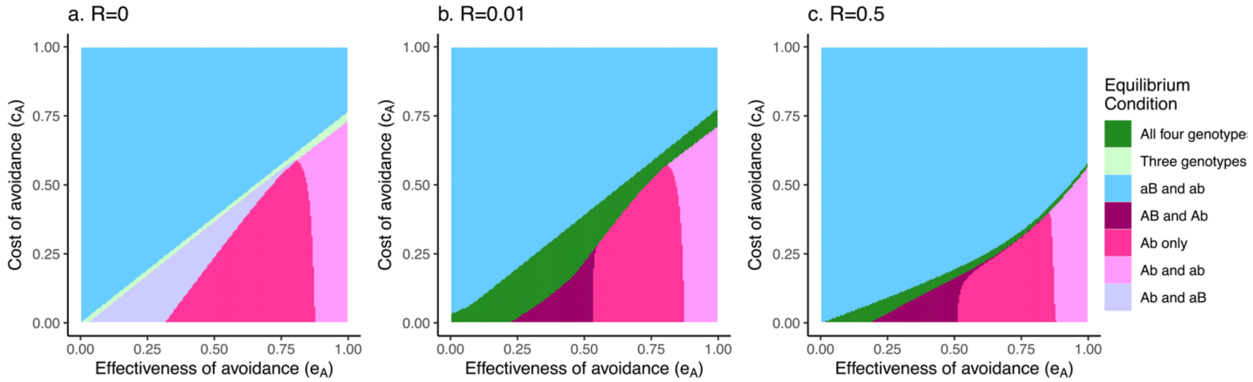


**Figure S6. Evolution of avoidance as $Ab$ from a low frequency under conditions that supported resistance polymorphism (Point III).** The cost and effectiveness of resistance were set at the values represented by Point III in Figure 1 ($e_B = 0.85, c_B = 0.65$) and run for 2000 timesteps, which allowed a polymorphism to be stably maintained between $B$ and $b$ when there was no variation in avoidance. Then $S_{Ab}$ was introduced at a low frequency ($10^{-5}$) and the simulations were continued for 2000 timesteps. Results varied with the recombination rate ($R$), and are illustrated in the panels (a. $R$ = 0, b. $R$ = 0.01, and c. $R$ = 0.5)

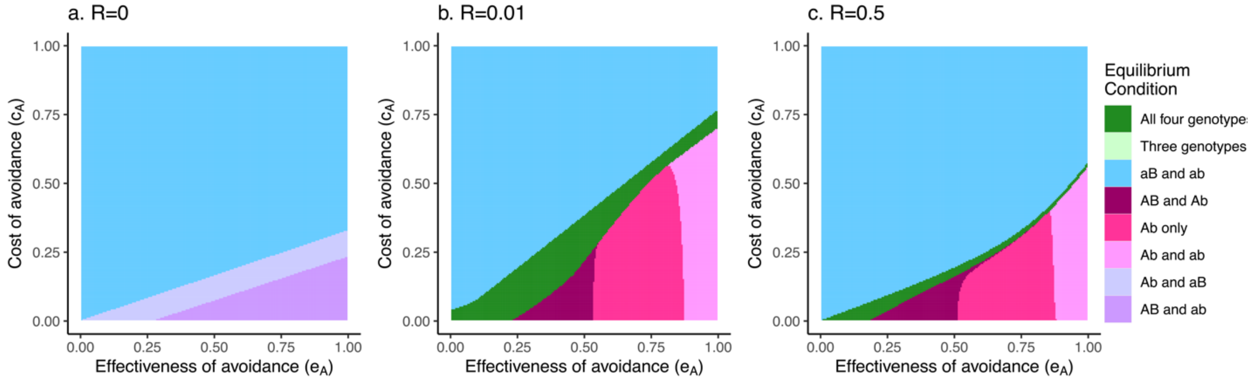


**Figure S7. Evolution of avoidance as $AB$ from a low frequency under conditions that supported resistance polymorphism (Point III).** The cost and effectiveness of resistance were set at the values represented by Point III in Figure 1 ($e_B = 0.85, c_B = 0.65$)and run for 2000 timesteps, which allowed a polymorphism to be stably maintained between $B$ and $b$ when there was no variation in avoidance. Then $S_{AB}$ was introduced at a low frequency ($10^{-5}$) and the simulations were continued for 2000 timesteps. Results varied with the recombination rate ($R$), and are illustrated in the panels (a. $R$ = 0, b. $R$ = 0.01, and c. $R$ = 0.5)
